# Saccadic network dysfunction underlies eye-movement variability in early Parkinson’s disease

**DOI:** 10.64898/2026.09.25.754307

**Authors:** Varsha Vasudevan, Sandy Mournet, Benoît Béranger, Caitlin Paige Rogers, Graziella Mangone, Isabelle Arnulf, Jean-Christophe Corvol, Marie Vidailhet, Stéphane Lehericy, Pierre Pouget, Nivethida Thirugnanasambandam, Cécile Gallea

## Abstract

Parkinson’s disease is a neurodegenerative disorder characterized by altered subcortical-cortical connectivity, leading to motor and cognitive deficits. Saccadic eye movement abnormalities are among the earliest and most persistent markers of the disease, serving as sensitive indicators of disrupted oculomotor and cognitive control. While changes in average saccadic behavioral measures are widely reported in Parkinson’s disease, intra-individual variability in saccadic performance and the underlying network dysfunction remain poorly understood. Here, we quantified intra-individual saccadic variability in early-stage Parkinson’s disease and identified its functional correlates within the resting-state saccadic network.

We analyzed data from 91 patients with early-stage Parkinson’s disease and 46 healthy controls who underwent oculomotor testing and resting-state functional magnetic resonance imaging. First, at the behavioral level, we assessed intra-individual variability in saccadic performance within each group using the coefficient of variation of reaction time and normalized peak velocity across trials. Second, at the brain level, we assessed group differences in static and dynamic functional connectivity across 39 regions comprising the saccadic network. We examined the relationship between network connectivity and saccadic variability using partial least squares analysis. We also quantified network organization using graph-theory measures of global efficiency and clustering coefficient. Finally, we tested the relevance of intra-individual saccadic variability and network features in distinguishing patients from healthy controls using a multivariate Mahalanobis classifier.

Our results showed that, compared to HC, patients had higher intra-individual variability in saccadic behavioral measures of reaction time (p_Holm_ = 0.012) and normalized peak velocity (p_Holm_ = 0.012). Interestingly, patients had (i) reduced static and dynamic connectivity within the saccadic network and (ii) weaker associations between network connectivity and intra-individual saccadic variability of specifically normalized peak velocity. These results were particularly driven by regions within the cognitive saccade network. The classifier that integrated intra-individual saccadic variability and network connectivity achieved 95.62% accuracy in distinguishing patients from healthy controls, with dynamic connectivity contributing more strongly to classification performance than static connectivity.

We demonstrate that elevated intra-individual variability of saccades, underpinned by altered functional connectivity within cognitive-oculomotor networks, is a prominent feature of early-stage Parkinson’s disease. Emerging as a promising physiological marker of disease pathophysiology, intra-individual variability offers a powerful new framework to refine early diagnosis and improve the tracking of neurodegeneration before severe clinical milestones are reached.

## Introduction

Parkinson’s disease (PD) is associated with widespread behavioral deficits spanning multiple domains, including gait, upper-limb coordination, speech, and oculomotor function. Among the earliest and most consistent manifestations of PD are abnormalities in saccadic eye movements, which can predict cognitive decline, gait disturbances, and overall disease progression^1–4^. Beyond mean performance deficits, PD is increasingly recognized as a disorder characterized by heightened intra-individual variability across motor domains^5–11^. These trial-to-trial fluctuations capture dimensions of motor control that average metrics fail to detect, offering a potentially more sensitive index of neurological dysfunction. In particular, intra-individual saccadic variability reflects subtle alterations within the oculomotor system^12–16^. Nevertheless, prior research in PD has overwhelmingly focused on mean performance, leaving trial-to-trial saccadic variability relatively underexplored. Consequently, how saccadic variability relates to underlying brain dynamics in PD remains poorly understood.

Saccade generation relies on a distributed brain network spanning cortical and subcortical regions^17–20^. Precise saccadic control requires synchronized interactions between prefrontal and parietal executive centers and subcortical motor circuits within the basal ganglia, brainstem, and cerebellum^21–23^. Given this distributed topology, the oculomotor system serves as a powerful model for examining how network dynamics govern behavior. While brain function was traditionally mapped using static connectivity, growing evidence underscores the importance of dynamic functional connectivity, i.e., transient reconfigurations in network topology that are uniquely vulnerable to pathological disruption^24–30^. Characterizing the dynamic properties of the saccadic network may therefore yield critical insights into the brain-behavior relationships that govern intra-individual saccadic variability.

Behaviorally, saccadic impairment in PD spans both motor and cognitive domains. Motor deficits encompass reduced accuracy of task performance, higher error rates, and altered kinematics (velocity, amplitude, and duration)^18,31–34^, whereas cognitive deficits manifest primarily as increased latencies in saccade planning and initiation^3,35–37^. Although cognitive impairments are well-established^1,36,38–40^, the heterogeneity of motor deficits^35,37,41,42^ highlights a major gap in our understanding of oculomotor pathophysiology. Given that saccade generation relies on the tight integration of cognitive and motor control^48–50^, these behavioral deficits likely arise from large-scale brain network reconfigurations^43–47^ within interconnected subcortical and cortical networks^51–53^. Dynamic interactions among cognitive and motor regions may be detrimental or could contribute to the maintenance of motor behavior^54,55^ and cognitive performance^30,43^. For instance, PD is characterized by functional reorganization across brain networks during choice tasks involving hand responses^33^, where interactions between cognitive and motor nodes may serve as compensatory mechanisms to counteract progressive neurodegeneration. Whether this functional reorganization extends to the saccadic network and directly accounts for eye-movement deficits in PD remains unknown. Specifically, how pathological shifts in dynamic connectivity of the saccadic network contribute to elevated intra-individual saccadic variability has not been established. Furthermore, the relative contributions of the cognitive and motor subdivisions of the saccadic network and their interactions to behavioral variability remain uncharacterized. Elucidating the relationship between dynamic network organization and oculomotor variability may therefore provide critical mechanistic insights into the neural architecture underlying saccadic dysfunction in PD.

The primary objective of this study was to investigate whether altered dynamics within the saccadic network, encompassing both its cognitive and motor sub-circuits, explain intra-individual variability in saccadic behavior in early PD. To test this, we enrolled patients with early-stage PD and age-matched healthy controls, quantifying both behavioral saccadic variability and intrinsic functional connectivity within the saccadic network using static and dynamic resting-state approaches. We tested three primary hypotheses: **(i)** patients with early-stage PD exhibit heightened intra-individual saccadic variability and altered functional connectivity within the saccadic network relative to healthy controls; **(ii)** the coupling between saccadic network connectivity and behavioral variability is significantly altered in PD; and **(iii)** because saccade generation relies on fluctuating, moment-to-moment network reconfigurations, dynamic functional connectivity should better characterize PD than static functional connectivity. We used a low-cognitive-load prosaccade task to evaluate saccadic variability. Although often termed a reflexive response, prosaccades depend on an elegant loop of spatial attention, target selection, and motor execution^20,21,23,56,57^. This methodological choice allowed us to isolate the neural interface between cognitive and motor networks without overloading the oculomotor system that is fragilized in PD patients. By integrating behavioral and neuroimaging metrics, this work advances our systems-level understanding of how cognitive and motor aspects of oculomotor control are disrupted in PD, highlighting its potential utility for early brain/behavior biomarker development.

## Materials and methods

### Participants and clinical data

The participants in this study were part of the monocentric prospective research cohort study conducted by the Paris Brain Institute (the ICEBERG study, ClinicalTrials.gov Identifier: NCT02305147). The cohort included healthy participants and individuals with PD. The patient inclusion criteria included (i) clinical diagnosis of PD (UK Brain Bank criteria) by sleep neurologists or movement disorder specialists, (ii) age between 18 and 75 years, (iii) no or subtle cognitive impairment (Mini-Mental State Examination score > 26/30), and (iv) time-from-diagnosis ≤ 4 years. Control participants were evaluated through clinical examination to verify the absence of a history of neurological or psychiatric disease. The study was approved by the local ethics committee (CPP Paris VI, RCB: 2014-A00725-42). All participants provided written informed consent, and the research was conducted in accordance with the Declaration of Helsinki. From the large cohort (see details in^58^), we included healthy controls (HC) and PD patients who underwent both MRI and oculomotor testing. Clinical assessment of patients was conducted using standardized scales, including the Movement Disorder Society Unified Parkinson’s Disease Rating Scale (MDS-UPDRS), Mini-Mental State Examination (MMSE), and Montreal Cognitive Assessment (MoCA). Oculomotor tests were performed at each visit under dopamine-ON conditions.

### Oculomotor data

#### Data acquisition and paradigm

Participants sat in a dark room with their eyes open in front of a 26-inch LCD monitor (ProLite Iiyama PL2600) at 60 cm (visual angle of 56° × 35°). Stimuli were displayed at 1920 × 1080 pixels and a 144 Hz monitor setting. Eye movements were recorded with a video-based eye tracker (Mobile EBT©, eyeBRAIN, France) at 500 Hz. Before the task, a standard nine-point calibration was conducted, requiring participants to fixate a 0.5° red target for 250 ms at each location. The spatial accuracy was approximately 0.25°-0.5°, and baseline noise was below 0.5°. The task was controlled using MeyeParadigm software, and eye-movement data were exported via MeyeAnalysis (v1.20.1).

The saccade task (**Supplementary Figure 1**) started with a central green fixation point (15 × 15 pixels; luminance 120 cd/m²) appearing on a black background (jittered interval of 2500– 3500 ms). After a 200 ms gap following the fixation period during which the screen is blank, a red target of the same size appeared at ±25° along the horizontal meridian. The target location was randomized to the left or right and visible for 1000 ms. Participants were instructed to make a rapid eye movement to the target, then return to the central fixation for the next trial. Each participant performed 48 horizontal saccades in two or four blocks.

#### Data analysis

Oculomotor data were analyzed using MATLAB scripts (R2023a, MathWorks, MA, USA). Horizontal and vertical eye position signals were segmented into trials using event markers corresponding to fixation onset and target onset. Trial boundaries were defined from fixation onset to the onset of the subsequent trial. All trials to both left and right targets were combined and analyzed. Position signals were smoothed using a Savitzky–Golay filter (order = 5, frame length = 21 samples). Trial-wise baseline drift was removed by subtracting the mean eye position computed over a 20-ms window immediately preceding target onset. Angular displacement was calculated from the drift-corrected horizontal and vertical components. Velocity was obtained using Savitzky–Golay differentiation kernels.

Saccades were detected using a velocity-based algorithm. For each trial, the peak velocity following target onset was identified, and saccade onset and offset were defined as the first and last time points at which velocity crossed 10% of the peak velocity before and after the peak, respectively. Reaction time was the interval between target onset and saccade onset. Trials were excluded if (a) saccade duration exceeded 200 ms, (b) signals showed saturation or flat-line artifacts, (c) baseline was unreliable, or (d) blink artifacts were present during saccade. For all valid trials, we extracted saccade amplitude, peak velocity, reaction time, and duration. Outliers were identified for each parameter using a ±3-standard-deviation criterion relative to the subject-specific mean and were excluded from further analyses. Trials were concatenated across blocks before subject-level analyses.

We quantified reaction time as a measure of saccadic decision-making and normalized peak velocity as a marker of saccadic kinematic change. For each participant, we computed the mean and variance across trials for reaction time, amplitude, peak velocity, and duration. To account for the main-sequence relationship between saccade amplitude, duration, and peak velocity, a subject-specific linear regression without intercept was fitted between amplitude and the product of peak velocity and duration^59^. The resulting slope parameter was used to generate predicted peak velocity values, and a normalized peak velocity (NormPV) was computed as a ratio of the observed to predicted peak velocity. The mean and variance of NormPV were then computed at the subject level. To quantify intra-individual saccadic variability relative to mean performance, the coefficient of variation (CV) was calculated as the standard deviation divided by the mean for reaction time and normalized peak velocity (CV-RT and CV-NormPV, respectively). These subject-level behavioral measures were used for subsequent group-level statistical analyses.

### Neuroimaging data

#### Data acquisition

MRI data were acquired for all participants using a 3T Siemens Prisma scanner (Siemens Healthineers, Erlangen, Germany) equipped with a 64-channel head coil. High-resolution structural images were obtained using a T1-weighted three-dimensional MP2RAGE sequence (TR = 5000 ms, TE = 2.98 ms, voxel size = 1 × 1 × 1 mm³). Resting-state functional images were collected using a multiband single-shot gradient-echo echo-planar imaging sequence (TR = 2190 ms, TE = 25 ms, flip angle = 90°, voxel size = 2.8 mm isotropic, 44 interleaved slices, multiband acceleration factor = 210 volumes). In addition, a shortblipped EPI acquisition (6 volumes) with reversed phase encoding was acquired to enable geometric distortion correction.

#### Data analysis

Functional MRI preprocessing was performed using Statistical Parametric Mapping (SPM12; Wellcome Trust Center for Neuroimaging, London, UK) implemented in MATLAB. Preprocessing steps included distortion and slice-timing correction, realignment for head-motion correction, artifact detection (ART-based identification with outlier scrubbing), spatial normalization to the Montreal Neurological Institute (MNI) template using DARTEL, and spatial smoothing with a 6-mm full-width-at-half-maximum Gaussian kernel.

Residual BOLD volumes were obtained by fitting a general linear model with nuisance regressors including white matter and cerebrospinal fluid signals, 24 motion parameters (six rigid-body motion parameters and their temporal derivatives and squared terms), and stick regressors identifying volumes with framewise displacement exceeding 0.5 mm and physiological confounds to write the residuals as cleaned 4D volumes. These cleaned volumes were then temporally band-pass filtered (0.01–0.1 Hz) using a fast Fourier transform–based approach to remove low-frequency drift and high-frequency physiological noise.

Region-of-interest (ROI) time series were extracted using custom MATLAB scripts built on SPM12. A predefined saccade network comprising 39 ROIs (**Supplementary Table 1**) was defined based on prior studies and meta-analyses examining the cortical and subcortical networks underlying pro-saccade generation^19,60–68^. ROIs encompassed frontal and parietal cortices, basal ganglia, cerebellar, brainstem, and thalamic structures. The ROIs were defined as described in the Supplementary Methods (subsection -Regions of Interest). Each ROI mask in MNI space was resliced to the functional image space, and the representative time series for each ROI was computed as the first eigenvariate of the voxel time series within the ROI mask using singular value decomposition, capturing the dominant pattern of coherent BOLD activity in the region.

The saccade network (defined using 39 ROIs) was subdivided using an operational functional classification into cognitive ROIs (ROIs 1–15 in **Supplementary Table 1**) comprising regions implicated in saccadic decision-making and higher-order control, and motor ROIs (ROIs 16– 39 in **Supplementary Table 1**) comprising regions primarily involved in oculomotor execution and kinematic control. This operational division was based solely on functional roles described in prior studies^57,69–73^.

#### Functional connectivity

We computed the magnitude of communication between two nodes, i.e., their weighted statistical synchronization using Pearson’s correlation. We computed the dynamic connectivity matrix for each participant using a sliding-window approach^74^. A window length of 120 seconds was used (window size = round (120/TR) = 55 time points; TR = 2.19 s). Although longer than the typically used windows, this choice ensured reliable sampling of low-frequency dynamics and reduced spurious variability in sliding-window correlations, based on theoretical considerations of signal reliability^75^. We analyzed 220 time windows, each including 55 time points and centered on the middle time point (symmetric window). At scan boundaries, windows were zero-padded to preserve a constant window length. Within each window, pairwise Pearson correlations among all 39 ROI time series were computed, resulting in 1521 edges (derived from 39 ROIs; 39 × 39 = 1521). The analysis yielded a time-resolved 3D connectivity matrix for each subject (39*39*220), with one correlation value per connection at each time window. The dynamic connectivity matrices were Fisher-z transformed. For dynamic connectivity (DC), the Fisher z-transformation was applied at each time window. The matrices were subsequently thresholded using a permutation-based null model derived from temporally shuffled time series^76,77^. For each subject, ROI time series were independently permuted along the temporal dimension, preserving amplitude distributions while disrupting temporal dependencies. To construct the null model, ROI time series were shuffled independently for each region and permutation (1000 iterations per subject). A correlation matrix was computed for each permuted dataset, and the resulting matrices were Fisher-z-transformed. The resulting Fisher-z-transformed coefficients were aggregated across permutations to generate a subject-specific null distribution of connectivity strengths. A pooled null distribution was then obtained by combining values across all subjects. The statistical threshold was defined as the 95th percentile of this pooled null distribution. Weighted adjacency matrices were constructed by retaining suprathreshold Fisher z-values and setting subthreshold connections to zero. For time-resolved DC, the same threshold was applied at each time window to derive dynamic adjacency matrices.

The functional connectivity matrix was partitioned into within- and between-network components based on the anatomical classification of the ROIs. Connectivity among the cognitive ROIs (ROIs 1-15) was defined as the cognitive saccade network (CSN), whereas connectivity among the motor ROIs (ROIs 16-39) was defined as the motor saccade network (MSN). Connectivity between the cognitive and motor ROIs was defined as the cognitive-motor saccade network (CMSN). Thus, CSN and MSN represent intra-network connectivity, while the CMSN represents inter-network connectivity (see **Supplementary Figure 2**).

The computation, thresholding, and analysis of static functional connectivity are described in the Supplementary Methods (subsection - Static connectivity).

#### Saccadic network-behavior coupling

To quantify the contribution of saccadic network connectivity to behavioral variability, we performed partial least squares regression using functional connectivity measures as predictors and behavioral variability measures as responses. Separate analyses were conducted to evaluate the relative contributions of intra-network (CSN and MSN) and inter-network (CMSN) connectivity. We also assessed whether these saccadic network-behavior relationships differed between HC and PD groups.

We considered DC network measures to investigate the contribution of the saccadic network in explaining the behavioral variability of saccades. Network-behavior coupling was examined using partial least squares (PLS) regression. Predictor variables in the network comprised the vector of correlation coefficients from the upper triangle of the connectivity matrix (n=741, since it was symmetric) for all subjects in a group, and the behavioral variables CV-NormPV and CV-RT for the subjects. Separate models were estimated for the HC and PD groups. The regression coefficients obtained from the fitted models were used to quantify the strength and direction of coupling between the network connectivity and saccadic variability. These saccadic network-behavior coupling values were then compared between HC and PD to assess disease-related alterations in the saccadic network’s contribution to behavioral variability.

##### Contribution of the cognitive and motor saccade networks

To determine the relative contributions of CSN, MSN, and CMSN, PLS regression was independently refit for each network, using only the edges within that network as predictors. The resulting network-specific connectivity-behavior coupling coefficients were then compared between HC and PD to assess disease-related alterations in each network’s contribution to saccadic behavioral variability.

The analogous method for static functional connectivity is described in the Supplementary Methods (subsection - Static connectivity).

#### Network organization characterization

To quantify the saccadic network organization, we computed graph-theoretical measures using functions from the Brain Connectivity Toolbox (Version 2019-03-03). Executing a saccade requires an optimal balance between (i) local network segregation with specialized processing in distinct networks, which we captured by measuring the clustering coefficient, and (ii) global network integration with rapid data sharing across networks, which we captured by measuring the global efficiency. Before estimating clustering coefficients, the weight matrices were normalized to the range [0,1]. Analyses were performed separately for static connectivity (SC) and dynamic connectivity (DC) matrices without thresholding. Static graph metrics consisted of a single value per subject, derived from subject-specific weighted adjacency matrices, whereas dynamic graph metrics were computed on each time-resolved weighted adjacency matrix. The subject-level values for DC were defined as the mean of the metric computed across all time windows. All graph metrics were computed at the subject level and subsequently used for group-level statistical comparisons between HC and PD patients.

#### Multivariate classification of PD

We tested whether intrinsic resting-state functional connectivity and saccadic behavioral variability serve as biomarkers of early PD. If so, connectivity- and behavior-derived variables should reliably distinguish PD patients from HC. To test this, we performed multivariate classification using features that showed significant group differences in prior analyses.

We derived global network measures from the entire saccade network, including cognitive and motor components. We reduced connectivity matrices to single-subject-level measures representing overall connectivity strength. For static connectivity, the overall connectivity strength was calculated as the mean of all upper-triangular values of each subject’s SC matrix (SC-avgcorr). For dynamic connectivity, we calculated overall connectivity strength as the mean of the upper-triangular values of each subject’s dynamic mean connectivity matrix (DC-avgcorr).

Classification was performed using eight subject-level features: static and dynamic overall connectivity strength (SC-avgcorr, DC-avgcorr), static and dynamic clustering coefficients (SC-CC, DC-CC), static and dynamic global efficiencies (SC-Eff, DC-Eff), and behavioral variability measures (CV-NormPV, CV-RT). All features were z-scored across participants before analysis. Classification was implemented using a Mahalanobis distance–based nearest-centroid classifier^78,79^ which accounts for differences in feature variance and covariance and is therefore well suited to heterogeneous group dispersion. Group centroids and covariance matrices were estimated for each group, and each subject was assigned to the group yielding the smaller Mahalanobis distance. Performance was quantified using classification accuracy and receiver operating characteristic (ROC) analysis with area under the curve (AUC). Robustness was assessed using leave-one-out cross-validation (LOOCV), and feature contributions were evaluated via leave-one-feature-out ablation analyses, with LOOCV performance recomputed after each ablation.

#### Statistical analysis

All statistical analyses were performed in MATLAB (R2023a, MathWorks, MA, USA). Statistical significance was set at p < 0.05 (two-tailed) unless otherwise specified. For subject-level behavioral measures (reaction time and normalized peak velocity) and graph metrics (clustering coefficient and global efficiency from SC and DC matrices), normality was assessed using the Lilliefors test. Between-group differences (HC vs PD) were evaluated using independent-samples *t*-tests when both groups were normally distributed, and Wilcoxon rank-sum tests otherwise. Test statistics are reported as *t*-or *z*-values, as appropriate.

For group-level statistical comparisons of the connectivity matrices, we used non-parametric permutation testing. We averaged the weighted adjacency matrices across time windows to obtain the DC mean matrix for each subject. These subject-wise average connectivity measures were then used for group-level statistical comparisons. The upper-triangle values of the DC mean matrix of each subject were used. To calculate observed group differences, we combined these values across all ROIs for all subjects within a group (HC or PD) to form a group-specific distribution and then assessed statistical differences using the Wilcoxon rank-sum test. To obtain empirical p-values, group labels were randomly permuted 10000 times while preserving group sizes. For each permutation, the rank-sum statistic was recomputed, and the empirical P-value was calculated as the proportion of permuted statistics equal to or more extreme (95^th^ percentile) than the observed statistic. When no permutation exceeded the observed statistic, P was set to 1/10000.

In both the whole saccadic network and individual CSN, MSN, and CMSN analyses, we assessed within-group reliability of the coupling between network connectivity and saccadic behavior using bootstrap resampling. Between-group differences were tested using a subject-level group-label permutation of group labels, thereby accounting for the dependence among connectivity edges estimated from a single fitted model (for further details, see Supplementary Methods subsection - Dynamic network connectivity to saccadic behavior coupling). Holm-Bonferroni correction (p_Holm_) was applied within each family of behavioral comparisons to account for multiple comparisons.

## Results

### Participants and clinical data

Demographics for the 46 HC and 91 PD patients are shown in **Table 1**. The patient and healthy groups were age-matched, with an imbalanced gender ratio representative of the higher prevalence of PD in males ^80,81^. Other than MMSE, which showed a weak trend toward lower scores in PD, and saccade amplitude, which was significantly reduced in PD, no other demographic, cognitive, or saccadic metric differed between groups.

**Table 1:** Clinical demographics and average metrics of saccadic behavior.

|  | HC (n=46) | PD (n=91) |
| --- | --- | --- |
| Age (mean ± SD<sup>#</sup>, years) | 60.9 ± 9.3 | 62.38 ± 9.5 |
| Sex: male/female (ratio) | 22/24 (0.9) | 56/35 (1.6) |
| Disease duration (years) | - | 1.22 ± 1.0 |
| Levodopa dose equivalent | - | 300.0 ± 125.6 |
| Hoehn and Yahr score | - | 2.0 ± 0.2 |
| UPDRS III – “best-on” condition | - | 27.0 ± 7.1 |
| <sup>##</sup>MMSE (0-30) | **29.4 ± 0.9** | **29.0 ± 1.0** |
| MoCA (0-30) | 28.5 ± 2.0 | 28.0 ± 2.0 |
| Saccade reaction time (ms) | 234.6 ± 32.2 | 228.3 ± 32.1 |
| <sup>###</sup>Saccade amplitude (°) | **21.9 ± 1.0** | **20.7 ± 1.4** |
| Saccade duration (ms) | 76.5 ± 11.5 | 74.9 ± 10.9 |
| Saccade peak velocity (°/s) | 457.4 ± 65.0 | 447.6 ± 55.7 |

<sup>#</sup>SD: standard deviation; **significant differences are shown in bold**; <sup>##</sup>ranksum, $z = 1.98$ , $p = 0.048$ ; <sup>###</sup>ranksum, $z = 4.86$ , $p < 0.001$ .

### Saccadic behavior changes with PD

Saccadic behavior, quantified as CV-NormPV and CV-RT, combined measures of mean and variance in normalized peak velocity and reaction time, respectively (as described in the Methods), differed significantly between PD and HC. CV-NormPV, which captured the combined effect of amplitude, duration, and peak velocity, increased for PD patients (data distribution: non-normal, rank-sum: z = -2.51, p = 0.012, p_Holm_ = 0.012, **Figure 1A**). The CV was higher for reaction time also in PD patients (data distribution: normal, t-test: t = -2.81, p = 0.006, p_Holm_ = 0.012, **Figure 1B**). Separate analysis of the differences in mean and variance showed that the CV difference was driven by the significant difference in variance between HC and PD for both normalized peak velocity (rank-sum: z = -2.53, p = 0.011) and reaction time (rank-sum: z = -2.25, p = 0.024). There was no significant difference in the means of normalized peak velocity (rank-sum: z = -0.52, p = 0.605) or reaction time (t-test: t = 1.10, p = 0.274) between the groups. Interestingly, the variability was greater in magnitude for reaction time than for normalized peak velocity.

**Figure 1.**
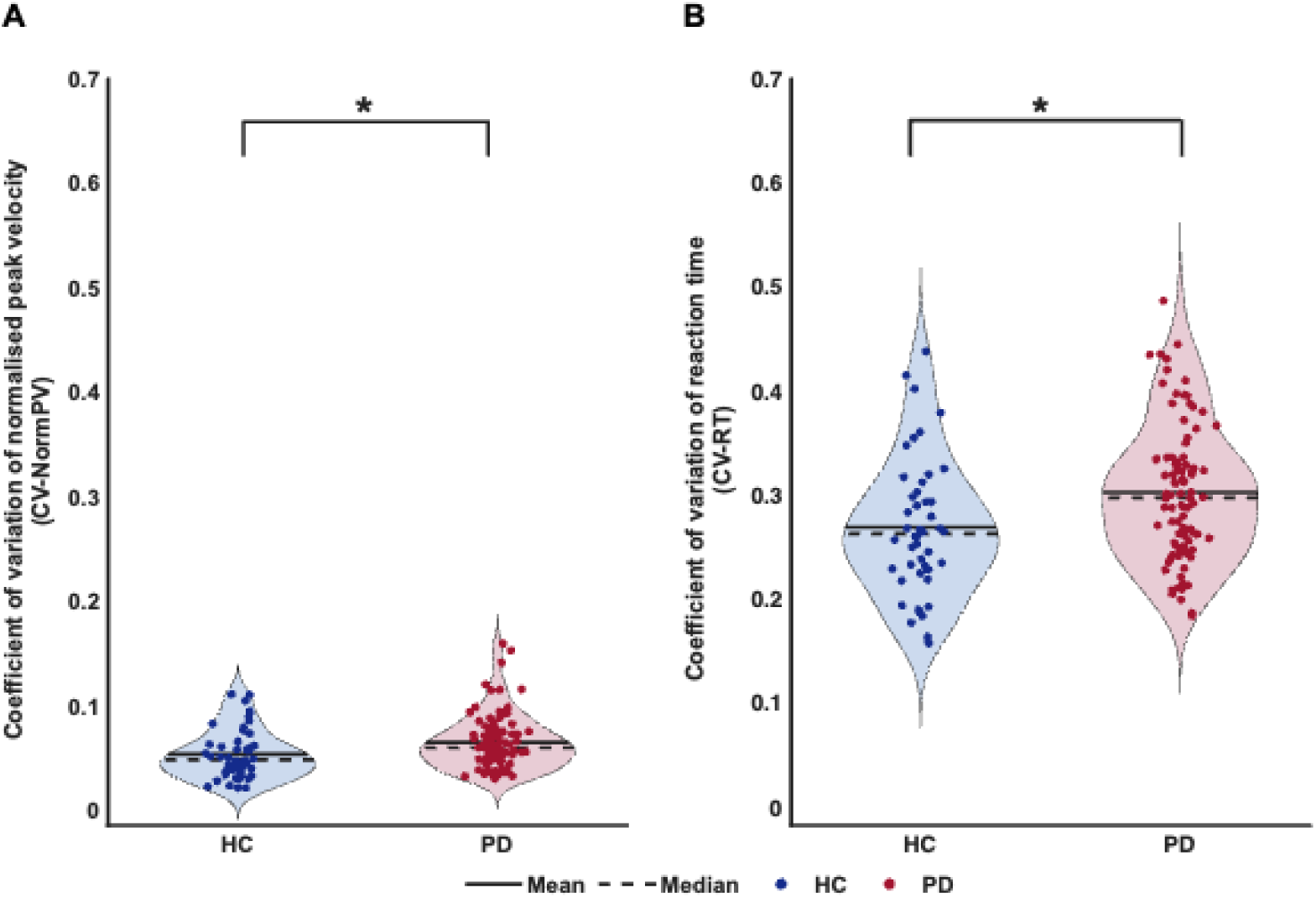
Group difference in the variability of saccadic behavioral parameters. Violin plots show the distribution of Coefficient of Variation (CV) values across participants in each group. **A.** CVs of normalized peak velocity, CV-NormPV. **B.** CVs of reaction time, CV-RT. Data are displayed in blue for healthy controls and in red for PD patients. The width of each violin reflects the density of observations, while individual points represent observations from individual participants. Solid and dashed horizontal lines denote the mean and median, respectively. Statistical thresholds for significant group differences are indicated by an asterisk (*p_Holm_ < 0.05).

### Saccadic network connectivity changes in PD

Both SC and DC were lower in PD, as shown by the group-averaged global network measure, i.e., the average connectivity across all edges of the connectivity matrix of the whole saccadic network (**Figure 2A**). The edgewise difference in the mean DC calculated across the time windows revealed a distributed pattern across the saccadic network ROIs in the group differences obtained using a non-parametric label-permutation approach (n = 10000, 95th percentile threshold = 0.0314). Of the total 741 edges in the upper half of the connectivity matrix excluding the diagonal, 515 edges (69.5%) showed significant differences, most of which (99.8%) had reduced DC in PD (**Figure 2B**). SC also yielded similar group differences (**Supplementary Figure 3**). Of the 515 edges that showed significantly reduced connectivity in PD, the largest proportion involved the CSN (42.5%), followed by the MSN (28.2%) and CMSN (29.2%). Only 1.3% of altered edges showed increased connectivity in the PD group, primarily in the MSN. These findings indicate a widespread reduction in connectivity across the saccadic network in PD, affecting all three functional subnetworks, with the largest proportion of altered connections involving the CSN.

**Figure 2.**
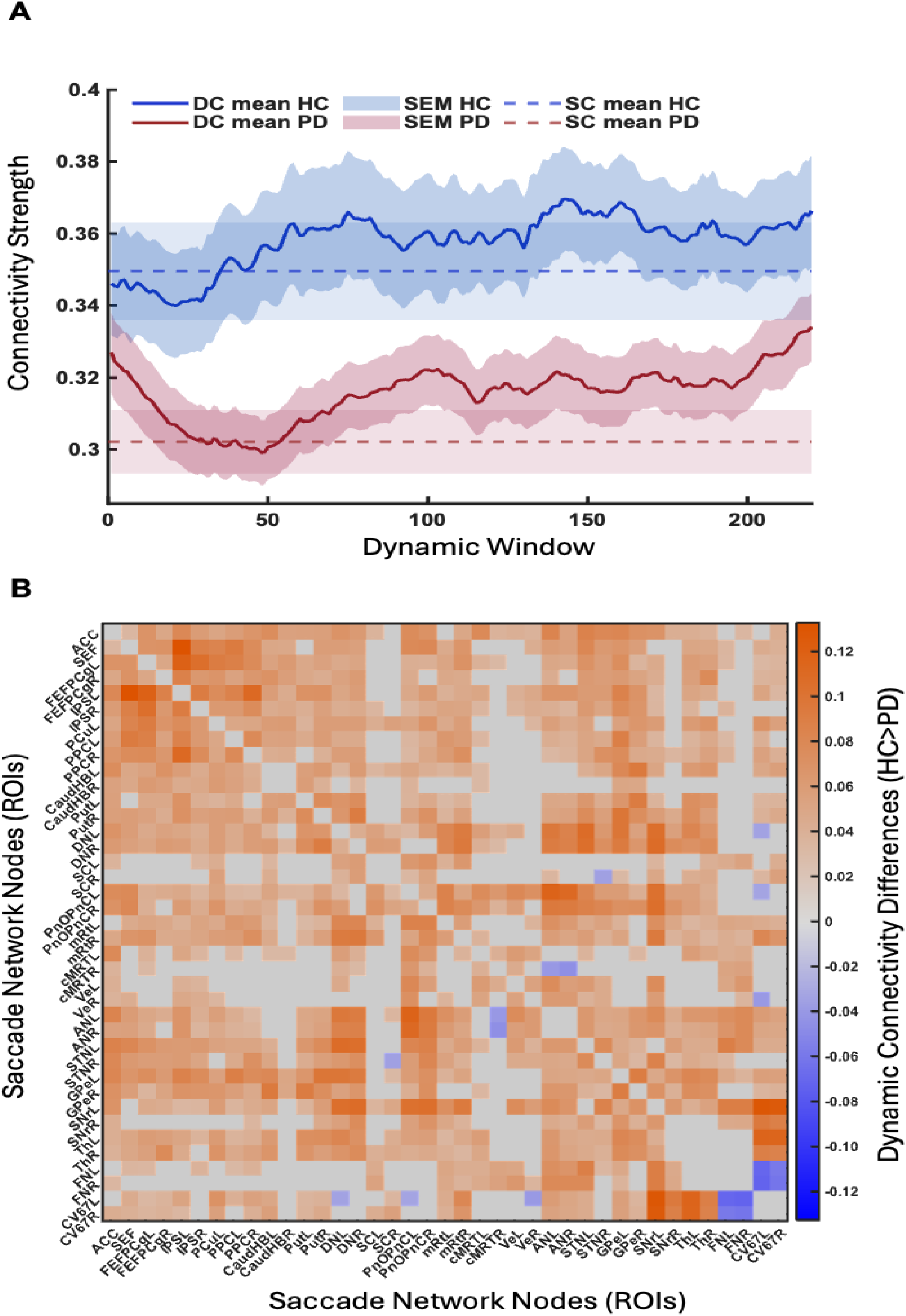
Group differences in functional connectivity. **A**. Static connectivity (SC; dashed lines) and dynamic connectivity (DC; solid, bold lines) are shown on a single graph to highlight differences between groups: in PD, both static and dynamic connectivity are lower than in HC. **B.** Dynamic connectivity matrix displaying the spread of group differences over the edges. The color bar shows the actual differences in Pearson correlation coefficients, and the gray edges are the non-significant ones after permutation testing.

### Saccadic network - behavior coupling

The partial least squares regression analysis used 741 connectivity values from the upper half of the connectivity matrices to predict two behavioral measures (CV-NormPV and CV-RT) in HC and PD separately. Here, we highlight the results of analyses using DC measures. Analyses using SC measures yielded similar results (**Supplementary Figure 4**).

### Saccadic network contribution to intra-individual saccadic variability

In HC, the connectivity pattern of the saccadic network explained CV-NormPV and CV-RT differently from PD (**Figure 3A and 3B**, radial lines, blue - HC and red - PD). Higher connectivity was reliably associated with greater CV-NormPV (p=0.002, p_Holm_ = 0.005; **Figure 3C**, blue violin), while network connectivity was unrelated to CV-RT (p=0.77, p_Holm_= 0.77; **Figure 3D**, blue violin). In contrast, network connectivity in PD patients showed no reliable relationship to either measure of saccadic variability (CV-NormPV: p=0.49, p_Holm_ = 0.54; CV-RT: p = 0.54, p_Holm_ = 0.54; **Figure 3C-D**, red violins). Comparing PD and HC, mean network-behavior coupling differed significantly between groups for CV-NormPV (p=0.01, p_Holm_ = 0.02), but not for CV-RT (p=0.17, p_Holm_ = 0.17). Full statistical details for the within-group and between-group analyses are provided in **Supplementary Table 2**.

**Figure 3.**
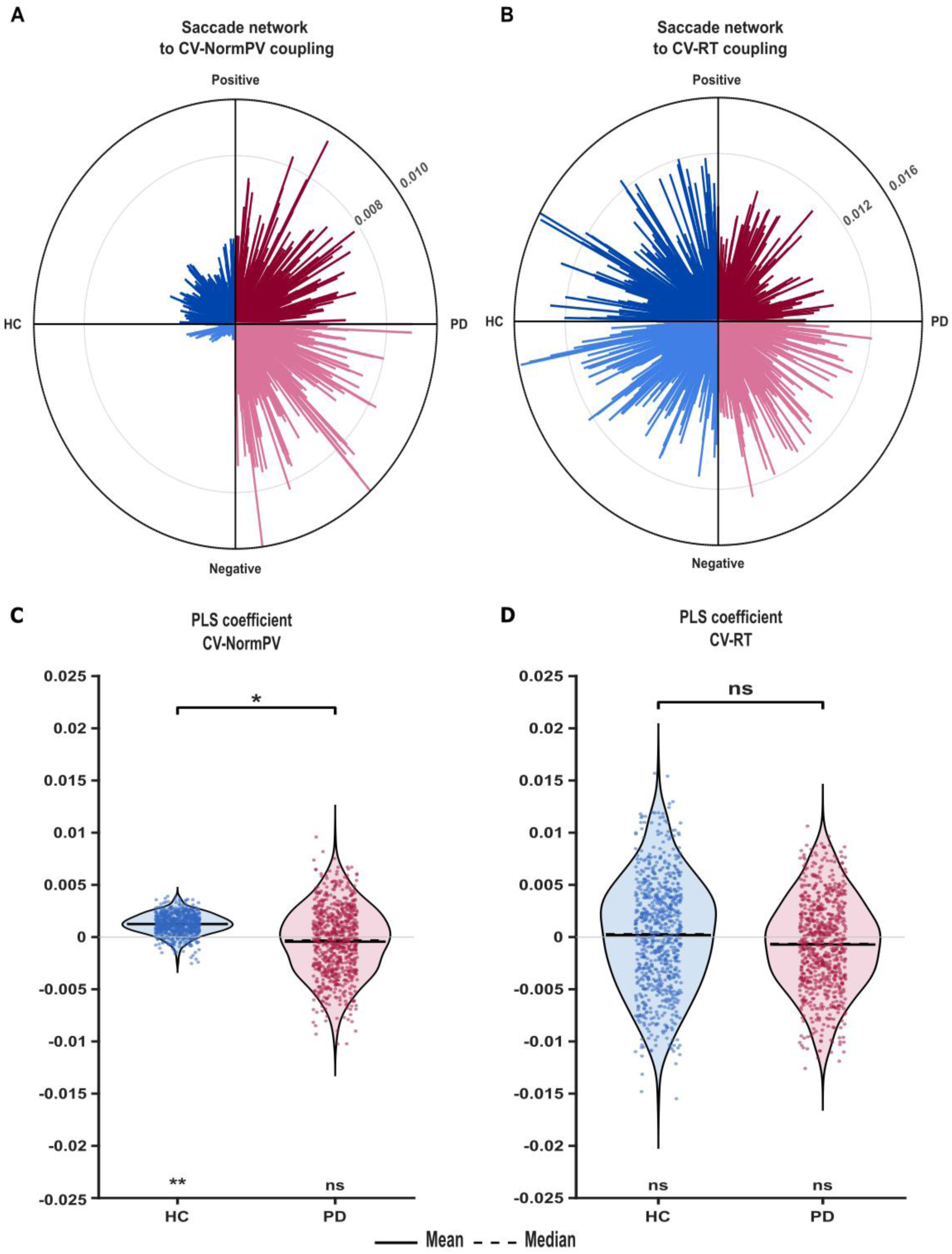
Coupling between dynamic connectivity in the saccadic network and saccadic variability. Network-behavior coupling was altered in PD compared to HC. **A-B**. Radial maps depict the distribution of network-behavior coupling coefficients relating dynamic connectivity to saccadic variability: CV-NormPV (**A**) and CV-RT (**B**), with the values denoted next to each. A-B. Radial line length represents the magnitude of the PLS regression coefficient, with positive and negative coefficients shown in the upper and lower quadrants, respectively. Blue and red represent HC and PD, respectively. **C-D**. Distributions of coupling coefficients between each edge (n=741) and saccadic variability in HC (blue) and PD (red) for CV-NormPV (**C**) and CV-RT (**D**). Positive coefficients indicate that higher connectivity was associated with greater saccadic variability, and vice versa. Violin plots show the distribution of coefficients across edges (represented by points); solid and dashed lines represent the mean and median, respectively. Statistical significance is indicated at the top of the graph for between-group comparisons and at the bottom for within-group significance relative to zero (*p_Holm_ < 0.05, **p_Holm_ < 0.01; ns, not significant).

### Contribution of cognitive and motor saccadic networks and their interactions to intra-individual saccadic variability

In HC and PD, network-behavior coupling followed different trends across all three saccade networks of CSN, MSN, and CMSN (**Figure 4A-C**).

**Figure 4.**
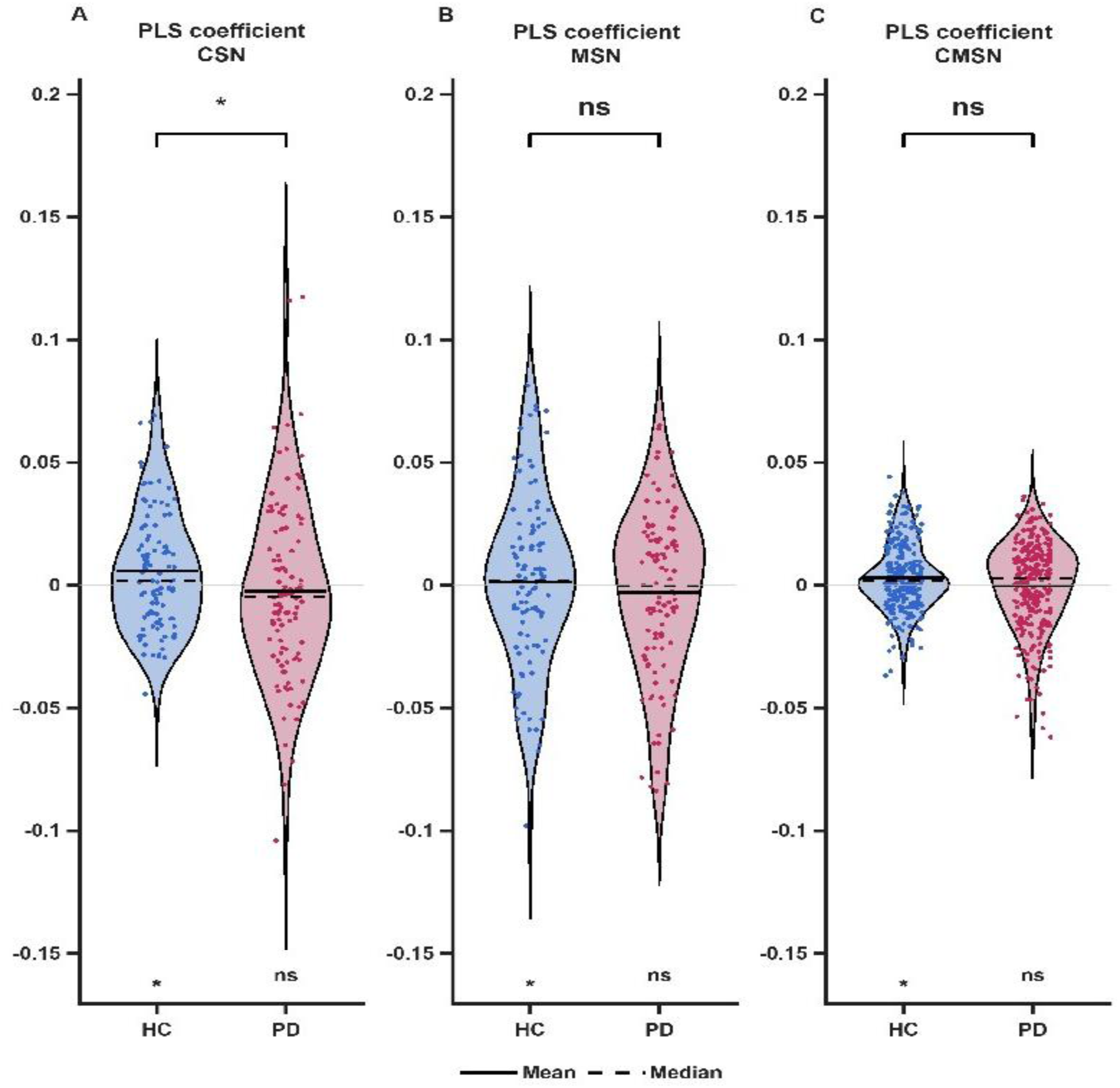
Specific contributions of cognitive and motor saccadic networks to saccadic behavior (CV-NormPV). Saccadic network-behavior coupling within and between the cognitive and motor networks was different between HC and PD, with significant differences in only the CSN. Violin plots show the distributions of coupling coefficients between network connectivity and saccadic variability (CV-NormPV) for the CSN (**A**), MSN (**B**), and CMSN (**C**) derived using a PLS regression model for each network. Blue and red colors represent HC and PD, respectively. Positive coefficients indicate that higher connectivity was associated with greater saccadic variability and vice versa. The plots depict the distribution of coupling coefficients across network edges (represented by points); solid and dashed horizontal lines represent the mean and median, respectively. Statistical significance is indicated at the top of the graph for between-group comparisons and at the bottom for within-group significance relative to zero (*p_Holm_ < 0.05, **p_Holm_ < 0.01; ns, not significant).

Within HC, connectivity showed a significant coupling to CV-NormPV in CSN (bootstrap resampling, p = 0.02, p_Holm_ = 0.02), MSN (bootstrap resampling, p = 0.006, p_Holm_ = 0.017), and CMSN (bootstrap resampling, p = 0.006, p_Holm_ = 0.017). This coupling was disrupted in PD, with no network connectivity showing a reliable relationship to CV-NormPV (CSN: p = 0.19, p_Holm_ = 0.59; MSN: p = 0.81, p_Holm_ = 1.00; CMSN: p = 0.76, p_Holm_ = 1.00). Comparing HC and PD directly, the between-group difference in coupling to CV-NormPV was significant only for CSN (group-label permutation, p = 0.006, p_Holm_ = 0.017) and did not reach significance for MSN (p = 0.07, p_Holm_ = 0.07). For CMSN, although there was a significant trend, this did not survive correction for multiple comparisons (p = 0.03, p_Holm_ = 0.06). For CV-RT, none of the within-group or between-group comparisons were significant (all p_Holm_ ≥ 0.37). The detailed statistics for all comparisons are presented in **Supplementary Table 3**.

Overall, saccadic network connectivity showed a reliable relationship with peak-velocity variability in HC that was absent in PD, both at the whole-network level and consistently across all three networks of CSN, MSN, and CMSN. The cognitive saccade network (CSN) showed the only significant between-group difference. No comparable relationship was detected for reaction-time variability at any level of analysis in either group.

### Network organization changes with PD

The reorganization of saccadic network topology was quantified using graph metrics. Significant group differences were observed in metrics derived from both SC and mean DC (**Figure 5A-D**). The clustering coefficient was lower in PD for both SC and DC connectivity (SC: t-test, t = 2.85, p = 0.005, p_Holm_ = 0.012; DC: rank-sum, z = 2.76, p = 0.006, p_Holm_ = 0.012; **Figure 5A, C**). Similarly, PD patients had reduced global efficiency (SC: t-test, t = 2.94, p = 0.004, p_Holm_ = 0.012; DC: t-test, t = 3.07, p = 0.003, p_Holm_ = 0.010; **Figure 5B, D**).

**Figure 5.**
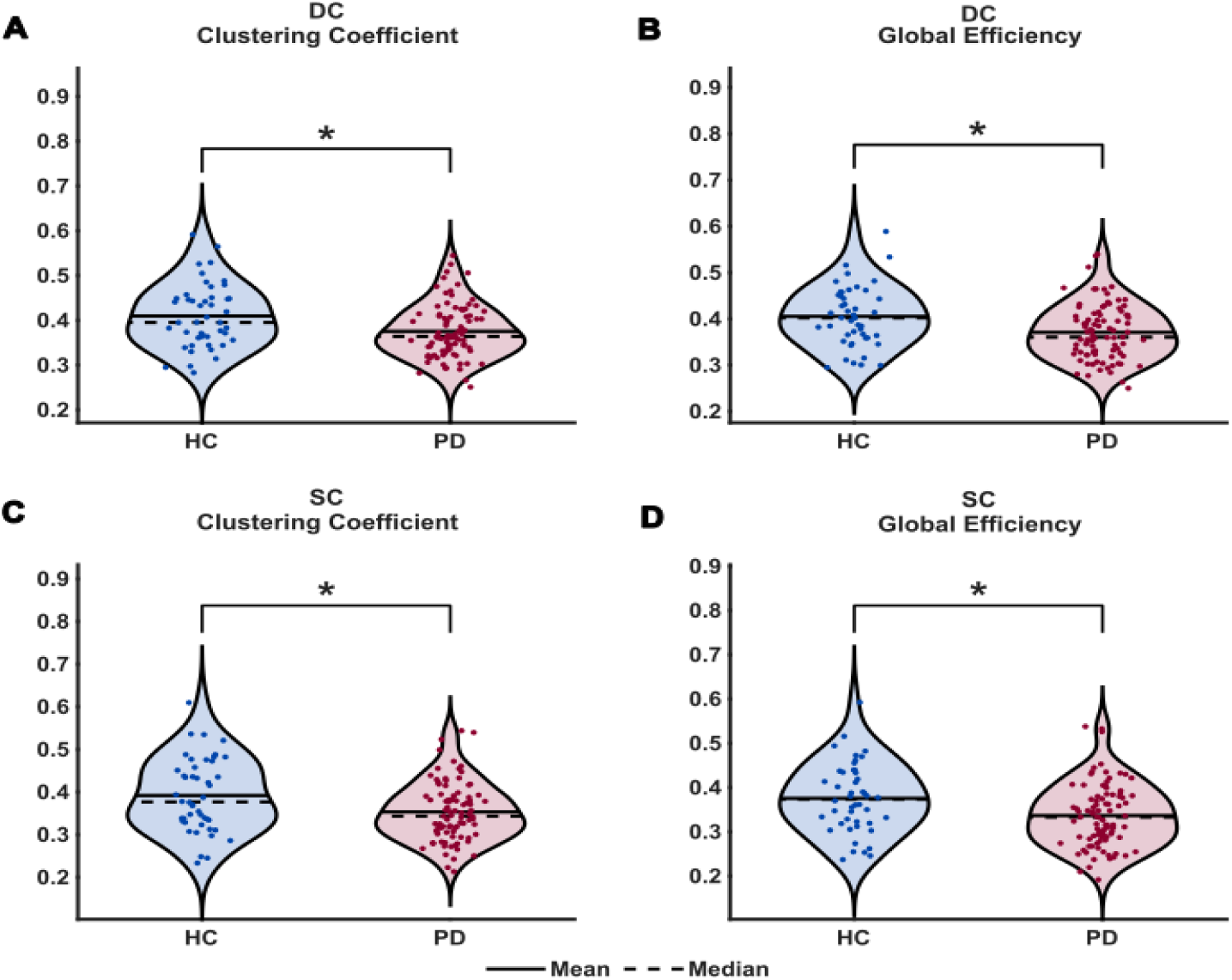
Network organization changes with PD. Violin plots show group differences in graph-theoretical network measures. **A-B.** Dynamic connectivity (DC). **C-D** Static connectivity (SC). **A, C:** Clustering coefficient. **B, D:** Global efficiency (right column). Healthy Controls (HC) = blue; Parkinson’s disease patients (PD) = red. Individual points represent participants. Solid and dashed lines denote the mean and median, respectively. PD exhibited significantly lower clustering coefficient and global efficiency relative to HC across both DC and SC measures. Asterisks on the brackets at the top of the plots indicate between-group comparisons (*p_Holm_ < 0.05).

### Multivariate classification of Parkinson’s disease

The classifier built using all 8 features representing network connectivity strength and organization, as well as behavioral variability, achieved a high classification accuracy of 98.5%, with only 2 HC individuals misclassified as PD patients (**Figure 6A**). The LOOCV, used to assess model generalizability to unseen individuals, yielded a classification accuracy of 95.6%, indicating robust and stable group separability across subjects. ROC curves and the full model exhibited near-ceiling performance, with high true positive rates achieved at very low false positive rates, confirming robust separability in the multivariate feature space (see bold black ROC curve in **Figure 6B**). The ROC performance degraded the most upon the removal of DC-based averaged connectivity strength (DC-avgcorr; see the downward shift of the dashed blue ROC curve compared to the bold black curve in **Figure 6B**). The ablation analyses showed that classification accuracy decreased to 61.3% when only DC features were removed, and to 83.9% when only SC features were removed (**Figure 6C**). In summary, the multivariate feature set showed robust separation of PD and HC, with DC connectivity strength contributing more strongly to classification than SC connectivity.

**Figure 6.**
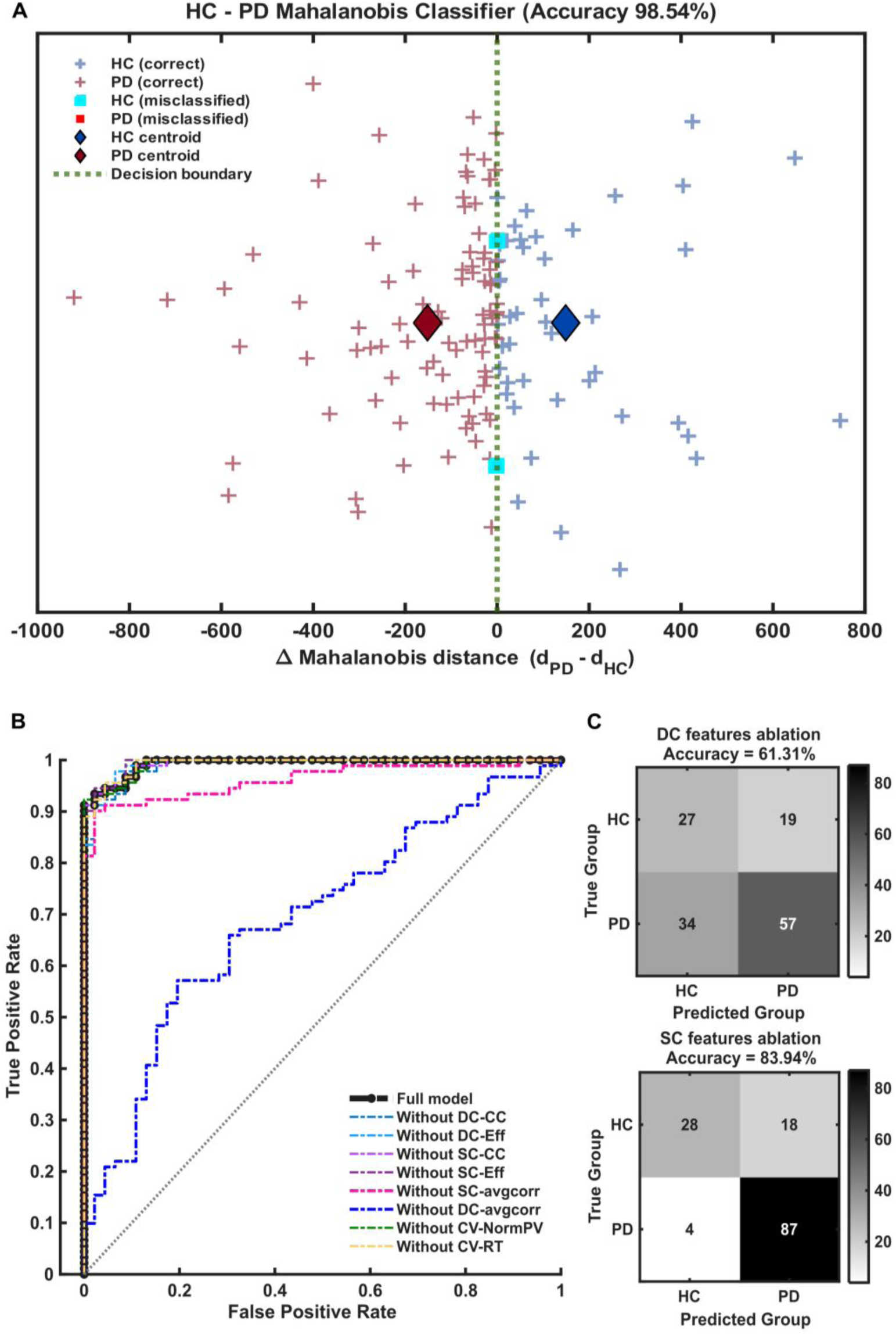
Classification of the population into healthy and PD groups using features derived from saccadic network connectivity and saccadic variability. **A.** Distribution of participants (each plus sign is a participant) over the manifold with HC (blue) and PD (red), showing misclassifications of two HCs (full cyan square) as PD, with an accuracy of 98.5%. Centroids of the groups are shown with filled diamonds. **B.** Classifier performance ROC curves with the full model in a bold black line, and an ablation analysis with each feature dropped, shown in different colors, showing that the ablation of the dynamic connectivity feature of the average correlation coefficient across ROIs has the maximum drop (blue dashed line). **C**. Leave-one-out classifier validation showing that the dynamic connectivity features ablation (lower panel), compared to static connectivity features (upper panel), has a greater effect on classification accuracy to separate PD from HC individuals.

## Discussion

In this study, we demonstrate that early-stage PD is characterized by a distinct threefold signature of elevated intra-individual saccadic variability, intrinsic saccadic network degradation, and decoupled brain-behavior interactions. Specifically, patients exhibited marked increases in saccadic variability accompanied by reduced functional connectivity within the saccadic network. Crucially, the breakdown in network-behavior coupling was primarily driven by functional reorganization within the cognitive areas primarily involved in oculomotor movement generation. At the topological level, graph-theoretical analyses revealed pervasive network disintegration, marked by diminished segregation and integration. Consequently, combining dynamic network properties with behavioral intra-individual variability robustly discriminates patients with PD from healthy controls, with dynamic connectivity features offering the strongest classification power. Collectively, these findings establish oculomotor intra-individual variability and dynamic network architecture as sensitive systems-level markers of early neurodegeneration.

Intra-individual variability represents a robust behavioral marker of saccadic impairment in early-stage PD. Our findings demonstrate that trial-to-trial saccadic variability may be a more sensitive feature explained by disrupted functional network connectivity, unlike mean saccadic performance, which may primarily reflect average neural activity and can therefore remain relatively preserved through compensation. These results support the emerging paradigm that behavioral variability reflects the stochastic dynamics of sensory, cognitive, and motor processing^82–84^ within intrinsically noisy neural systems. Rather than being dismissed as mere statistical noise, behavioral variability serves as an informative indicator of motor network integrity^85^. Within the oculomotor system, where kinematics are rapid and exquisitely quantifiable, intra-individual variability provides a sensitive window into central sensorimotor control^15,16^. Consistent with previous studies in PD^67,86–88^, patients exhibited marked increases in the variability of reaction time and peak velocity. Notably, the coefficient of variation for both reaction time and normalized peak velocity robustly differentiated patients with PD from healthy controls, even in the absence of significant differences in mean performance. Preserved mean performance may be maintained through transient compensatory mechanisms that obscure underlying network degradation, likely explaining the inconsistent findings in mean saccadic metrics reported across PD cohorts^89–92^. Similar increases in behavioral variability across distinct neurological disorders^93–96^ and effector systems^97–100^ suggest that heightened movement variability reflects a universal hallmark of pathological neural instability. Consequently, these findings reinforce the necessity of treating intra-individual variability not as residual noise, but as a robust, clinically meaningful dimension of neurodegenerative disease characterization.

Altered saccadic network dynamics represent a fundamental neurobiological hallmark of early-stage PD. In this study, we confirmed the hypothesis that PD is characterized by disrupted brain dynamics across interconnected neural systems. Although historically conceptualized primarily as a focal dopaminergic pathology, converging evidence now situates PD as a primary model of large-scale network dysregulation^86,101–105^. Our findings extend this network perspective to the oculomotor system, demonstrating that disease-related dysfunction reflects a widespread reorganization of cognitive and motor nodes across cortical and subcortical regions. While previous oculomotor research in PD has predominantly focused on regional activation differences or whole-brain alterations^50,62,106^, the primary novelty of the present work lies in characterizing how these specialized nodes interact as a coordinated system. Using functional connectivity and graph-theoretical metrics, we demonstrated pervasive reductions in saccadic network connectivity accompanied by decreased global efficiency and clustering coefficients. This pattern indicates a dual impairment in global integration and local segregation across the saccadic network, a system-level disintegration that remains undetectable when evaluating regional activity alone.

Whereas most resting-state functional connectivity studies in PD have relied on static correlations^102,107–110^, our findings highlight the critical added value of dynamic functional connectivity^111–114^. Dynamic metrics capture time-varying fluctuations in connectivity, serving as a proxy for network-level neural communication variability. This approach aligns with dynamical-systems frameworks of brain function, which posit that neural activity evolves over time within structured state spaces constrained by underlying anatomical architecture^115^. Within this framework, large-scale brain networks exhibit attractor landscapes that govern transitions between stable functional states; dynamic connectivity thus indexes how the brain traverses these energy landscapes^116^. In PD, the disruption of such dynamics manifests as (i) lower efficiency of communication reflecting reduced neural flexibility; and (ii) reduced clustering coefficient reflecting uncoupled coordination within the saccadic network nodes. Thus, this drives saccadic network-level disintegration and confers high sensitivity for disease classification. Future research quantifying transient network states, dwell times, and transition topologies in behaviorally grounded networks will be essential for uncovering the covert neural mechanisms driving subtle clinical manifestations.

Functional reorganization of the cognitive-motor network represents a hallmark of early-stage PD. By combining neuroimaging and oculomotor behavior in patients with early-stage PD, we demonstrate that disrupted functional connectivity directly underlies elevated intra-individual variability in saccadic behavior. In healthy controls, functional connectivity within cognitive and motor saccadic sub-networks was reliably associated with normalized peak velocity variability, while showing no association with reaction time variability, illustrating that behavioral stability of saccade kinematics depends on fine-tuned, coordinated network interactions. This aligns with prior literature describing decision-making and kinematic processes as tightly coupled within an integrated cognitive-motor architecture^117–120^. In PD, however, this coordinated brain-behavior topology was fundamentally altered, with network-behavior coupling to normalized peak velocity variability lost across the cognitive and motor networks and their interaction. This loss of coupling was robustly driven by the cognitive saccadic network, reflecting network-level disruption and partial functional reorganization. By decomposing the saccadic network into its constituent cognitive and motor sub-circuits, we uncovered the distinct contributions of these sub-networks to trial-to-trial behavioral variability, reinforcing the concept that behavioral variability is shaped by large-scale network architecture and is exquisitely sensitive to neurodegenerative pathology^121–123^.

In early-stage PD, functional alterations were most prominent within the cognitive saccadic network and extended across inter-network connections. This finding accords with the notion that cognitive control recruits distributed inter-network interactions (including recruitment of motor circuits)^124–126^ to support compensatory processes that preserve motor output despite progressive motor network degradation. Indeed, longitudinal neuroimaging evidence indicates that PD progression is shaped by dynamic cerebral compensation, wherein increased engagement of parieto-premotor circuits helps offset emerging motor deficits such as bradykinesia^127^. In this context, the altered coupling between saccadic variability and the cognitive sub-network (contrasted with the relative stability of the motor sub-network) could reflect an active compensatory mechanism. Collectively, these findings position cognitive-motor network reorganization as a pivotal systems-level bridging mechanism, linking altered neural dynamics to behavioral dysfunction in PD. Future studies could investigate whether non-invasive neuromodulation could serve as a targeted therapeutic tool to restore network-behavior coupling and rehabilitate motor stability.

Several methodological limitations of the present study warrant consideration. First, functional connectivity estimates were derived from resting-state functional MRI rather than task-based paradigms, which limits direct inference regarding online neural dynamics during active saccade generation. Second, all patients were evaluated in the dopaminergic ‘ON’ state; consequently, potential treatment-induced effects on both behavioral variability and network connectivity cannot be entirely ruled out. Third, although our machine-learning model achieved robust classification performance, external validation in larger, independent cohorts will be essential to establish the generalizability and clinical utility of eye-movement intra-individual variability and dynamic connectivity as robust markers of early PD. Finally, the observed imbalance in sex distribution between groups may have introduced subtle confounding influences on network topology and motor variability. Further studies utilizing larger, sex-stratified cohorts will be necessary to explicitly dissociate sex-specific effects from disease-related pathophysiological mechanisms.

In conclusion, this study establishes a clinical framework for brain–behavior integration in early-stage PD. Rather than reflecting isolated neural or behavioral deficits, PD manifests as a system-level disruption across interconnected networks. While separate individual behavioral or neuroimaging metrics provide incomplete clinical insights, their integration robustly distinguishes patients with PD from healthy controls. Saccadic eye movements represent an ideal translational tool: they are rapid, non-invasive, scalable, and probe distributed cortical– subcortical networks susceptible to neurodegenerative processes^56,128–131^. Combining saccadic variability with intrinsic network connectivity offers significant potential for early disease detection before overt clinical presentation, as well as for the differential diagnosis of atypical parkinsonian syndromes with distinct brainstem and cerebellar involvement^127^. Ultimately, because oculomotor dysfunction tracks key prognostic indicators (including freezing of gait and cognitive decline)^3,11,132–134^, behaviorally anchored network dynamics could establish a robust foundation for phenotyping, stratifying, and monitoring parkinsonian disorders.

## Data availability

The data supporting the findings of this study are not publicly available due to ethical and privacy restrictions involving human participant data. Aggregated data may be available from the corresponding author upon reasonable request and subject to institutional approval.

## Acknowledgements

This section is not mandatory.

## Supplementary material

### Supplementary methods

#### Saccadic task paradigm

The saccadic task has been described in detail in the main Methods section. Participants were instructed to initiate saccades starting from a fixation point to reach a lateral target, while eye movements were recorded. **Supplementary Figure 1** provides a schematic illustration of the experimental paradigm. Each trial started with a central fixation of jittered duration. The central fixation disappeared for 200 ms, followed by the appearance of one of two possible lateral targets, placed on the horizontal axis passing through the central fixation. Participants had 1000 ms to perform the saccade, after which the target disappeared.

**Supplementary Figure 1.**
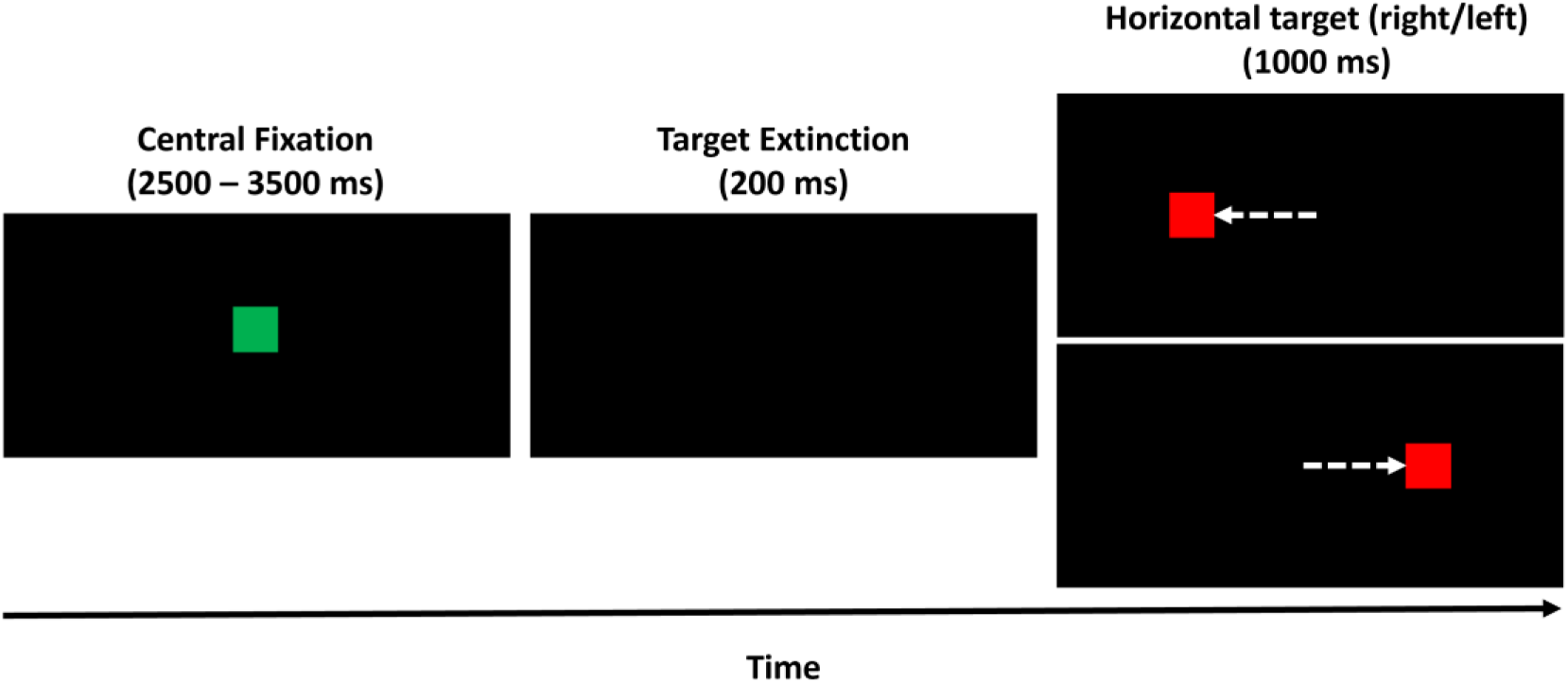
Saccade task paradigm. Illustration of a single trial timeline. The black rectangle represents the display screen. The green square denotes the central fixation stimulus, and the red square denotes the peripheral target stimulus. The white dashed arrow represents the instructed eye movement toward the target; this information is for illustrative purposes and was not seen by participants.

### Data analysis and statistics

#### Regions of interest

The ROIs were defined using a combination of functional (F) and anatomical (A) approaches to capture regions implicated in saccadic eye movements. The complete list of ROIs is provided in **Supplementary Table 1**, along with their abbreviations. The network comprises 15 cognitive and 24 motor ROIs. Functional ROIs were generated in MarsBaR (MARSeille Boîte À Région d’Intérêt), a toolbox for SPM using Montreal Neurological Institute (MNI) coordinates reported in previous neuroimaging studies of pro-saccade and anti-saccade tasks. For each reported activation peak, a spherical ROI with a radius of 6 mm was created around the corresponding MNI coordinate. Anatomical ROIs were obtained directly from the WFU_PickAtlas (WFUA, WFU_PickAtlas toolbox v3.0.5) or Brainstem Navigator Atlas (BSNA, Brainstem Navigator toolkit v0.9) using the corresponding atlas-defined anatomical masks. The anatomical atlas used for these ROIs is indicated in Supplementary Table 1. In cases where two functional coordinates were sufficiently close that their corresponding 6-mm spherical ROIs overlapped (e.g., the frontal eye field and precentral gyrus within the same hemisphere, or the left and right supplementary eye fields), the two ROIs were merged by creating a single 6-mm spherical ROI centered at the center of mass of the two coordinates. The coordinates of these merged ROIs are reported as the *Origin* in **Supplementary Table 1**. A 6-mm-radius sphere was then generated around this merged center coordinate.

**Supplementary Table 1:** Regions of interest defining the saccade network.

| ROI number | Region of Interest | Origin | Labels |
| --- | --- | --- | --- |
| **Cognitive ROIs/nodes saccade network** |  |  |  |
| **1** | Anterior cingulate cortex | F [6, 20, 44] | ACC |
| **2** | Supplementary eye field (combined left and right) | F [1, 5, 60] | SEF |
| **3** | Frontal eye field & precentral gyrus (left combined) | F [-40, -6, 50] | FEFPCgL |
| **4** | Frontal eye field & precentral gyrus (on right combined) | F [42, 0, 50] | FEFPCgR |
| **5** | Intra-parietal sulcus (parietal eye field) left | F [-24, -62, 40] | IPSL |
| **6** | Intra-parietal sulcus (parietal eye field) right | F [28, -52, 44] | IPSR |
| **7** | Precuneus left | F [-16, -68, 62] | PCuL |
| **8** | Posterior parietal cortex cluster left | F [-28, -54, 54] | PPCL |
| **9** | Posterior parietal cortex cluster right | F [22, -50, 54] | PPCR |
| **10** | Caudate head and body left |  | CaudHBL |
| 11 | Caudate head and body right | A (WFUA TD Brodmann Areas+) | CaudHBR |
| 12 | Putamen left |  | PutL |
| 13 | Putamen right |  | PutR |
| 14 | Dentate left |  | DNL |
| 15 | Dentate right |  | DNR |
| Motor ROIs/nodes of saccade network |  |  |  |
| 16 | Superior colliculus left | A (BSNA) | SCL |
| 17 | Superior colliculus right |  | SCR |
| 18 | Pontine reticular formation left | A (BSNA) | PnOPnCL |
| 19 | Pontine reticular formation right |  | PnOPnCR |
| 20 | Mesencephalic reticular formation left | A (BSNA) | mRtL |
| 21 | Mesencephalic reticular formation right |  | mRtR |
| 22 | Medullary reticular formation left | A (BSNA) | cMRTL |
| 23 | Medullary reticular formation right |  | cMRTR |
| 24 | Vestibular nuclei complex left | A (BSNA) | VeL |
| 25 | Vestibular nuclei complex right |  | VeR |
| 26 | Abducens nuclei left | A (BSNA) | ANL |
| 27 | Abducens nuclei right |  | ANR |
| 28 | Subthalamic nucleus left | A (BSNA) | 'STNL' |
| 29 | Subthalamic nucleus right |  | 'STNR' |
| 30 | Globus pallidus externa left | A (WFUA-TD Brodmann Areas+) | 'GPeL' |
| 31 | Globus pallidus externa right |  | 'GPeR' |
| 32 | Substantia nigra pars reticulata left | A(BSN) | 'SNrL' |
| 33 | Substantia nigra pars reticulata right |  | 'SNrR' |
| 34 | Thalamus left<br>(thalamoreticular, mediodorsal & ventrolateral nuclei) | A (WFUA-AAL) | ThL |
| 35 | Thalamus right<br>(thalamoreticular, mediodorsal & ventrolateral nuclei) |  | ThR |
| 36 | Fastigium oculomotor region left | A (WFUA-TD Labels) | FNL |
| 37 | Fastigium oculomotor region right |  | FNR |
| 38 | Cerebellar (oculomotor) vermis left | A (WFUA- AAL) | CV67L |
| 39 | Cerebellar (oculomotor) vermis right |  | CV67R |

#### Intra- and inter-network connectivity

The functional connectivity matrix was operationally divided into CSN and MSN, representing intra-network connectivity, and CMSN, representing inter-network connectivity, as discussed in the main methods section, Functional connectivity. **Supplementary Figure 2** illustrates the organization of the 39 × 39 connectivity matrix, resulting in 741 edges in the upper half of the symmetric matrix, and defines the CSN, MSN, and CMSN connectivity measures used throughout the analyses.

**Supplementary Figure 2.**
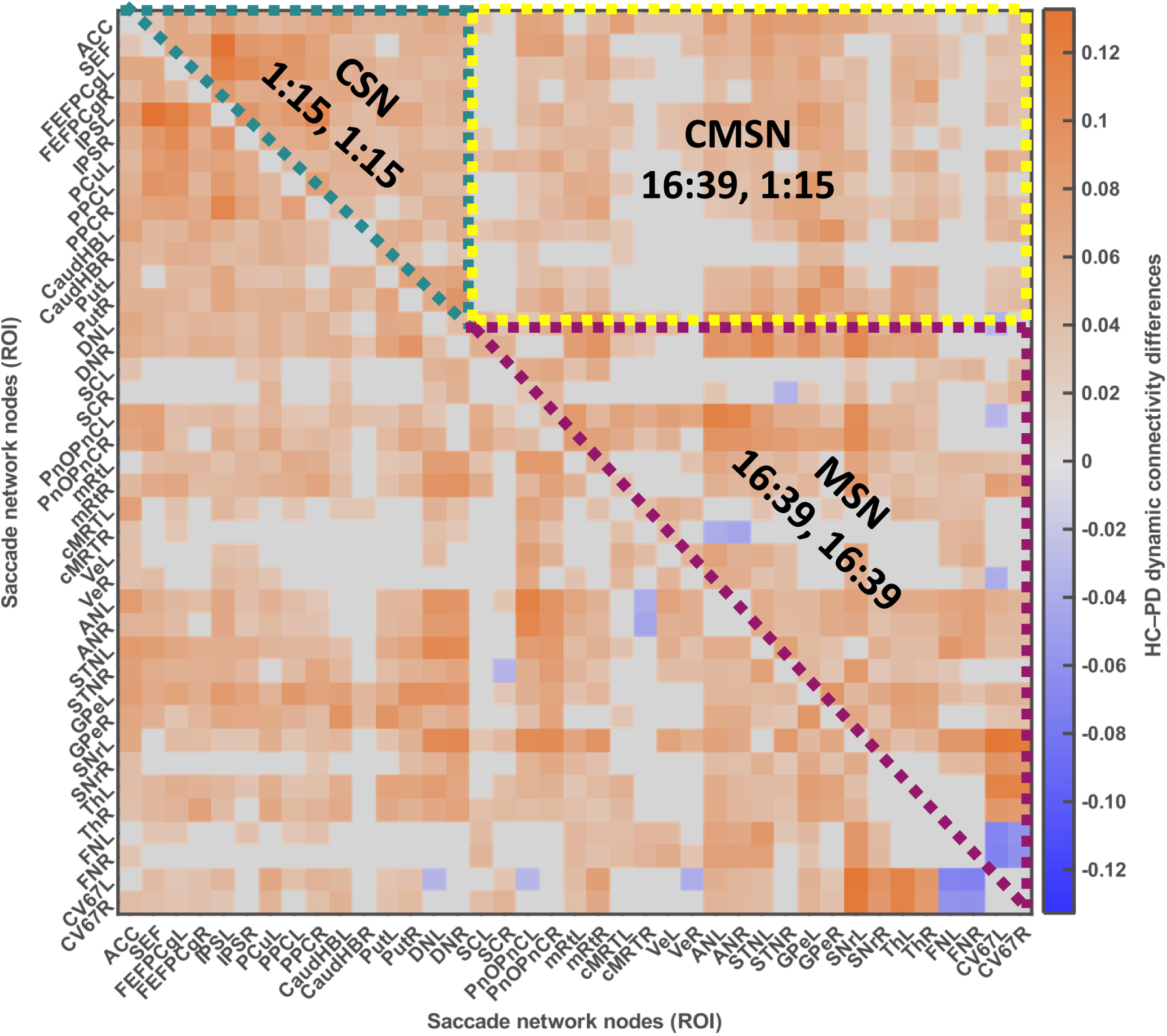
Definition of the Cognitive Saccade Network (CSN), Motor Saccade Network (MSN), and Cognitive–Motor Saccade Network (CMSN) within the functional connectivity matrix. ROIs 1–15 correspond to the cognitive, and ROIs 16–39 correspond to the motor subdivisions. The upper-left block (green) represents the CSN (intra-connectivity within cognitive), the lower-right block (purple) represents the MSN (intra-connectivity within motor), and the upper-right block (yellow) represents the CMSN (inter-connectivity between cognitive and motor connectivity). Owing to the symmetry of the connectivity matrix, only the upper triangular elements were retained for subsequent analyses.

#### Dynamic network connectivity and saccadic behavior coupling

Partial least squares (PLS) regression coefficients relating saccadic network connectivity to behavioral variability were derived separately for HC and PD. Because these coefficients are not independent observations, all edges within a group’s model are jointly constrained by the same small number of latent components from a single fit, two resampling-based procedures were used in place of conventional parametric or edge-level rank-based tests, which would treat non-independent edges as independent samples and inflate apparent significance.

##### Within-group reliability with bootstrap resampling

To test whether a group’s own mean network-behavior coupling reliably differed from zero, subjects within that group were resampled with replacement (5000 iterations); the PLS model was refit on each resample, and the mean coefficient across all edges was recorded. This produced an empirical distribution from which 95% confidence intervals were derived (2.5th–97.5th percentile). A group’s coupling was considered reliable if this interval excluded zero. A two-tailed p-value was derived from each bootstrap distribution based on the proportion of iterations falling on either side of zero.

##### Between-group comparison with subject-level permutation

To test whether mean coupling differed between HC and PD, group membership was randomly permuted across the pooled sample of 137 subjects (5000 iterations, preserving original group sizes of 46 HC and 91 PD). PLS models were refit separately on each shuffled grouping, and the resulting mean coefficient difference was compared to the observed HC-PD difference to derive a permutation p-value.

For both procedures, p-values were Holm-corrected within a family of two behavioral comparisons (CV-NormPV, CV-RT), applied separately to the within-HC, within-PD, and between-group test families. All analyses were implemented in MATLAB (plsregress).

##### Cognitive and motor network to saccadic behavior coupling

To assess the contribution of CSN, MSN, and CMSN specifically to saccadic behavior, a PLS regression model was refit separately with each of them, using only that sub-network’s edges as predictors, rather than extracting coefficients from the whole-network model. The same within-group bootstrap and between-group permutation procedures described above were applied to each sub-network, with Holm correction applied across the three sub-networks, computed separately for each behavior and group.

#### Static connectivity

Static functional connectivity was computed as the Pearson correlation between time series for all pairs of the 39 ROIs across the entire scan duration, yielding 1521 unique connectivity edges (39 × 39 = 1521). The resulting connectivity matrices were Fisher *z*-transformed and thresholded using the same permutation-based null model described for dynamic connectivity (main methods section, Dynamic Functional Connectivity). Briefly, a subject-specific null distribution was generated from 1000 temporal permutations of the ROI time series and pooled across subjects to define a statistical threshold (95th percentile). Weighted adjacency matrices were constructed by retaining suprathreshold Fisher *z* values and setting all remaining connections to zero.

Group differences in static functional connectivity were assessed using the same statistical framework as described in the main methods (Statistical Analysis). Briefly, subject-wise, weighted adjacency matrices were compared between HC and PD using non-parametric permutation testing (10,000 permutations) based on the upper-triangular connectivity values.

The contribution of static functional connectivity to saccadic behavioral variability was assessed using the same partial least squares regression framework, within-group bootstrap resampling, and between-group permutation testing procedures described in Supplementary Methods for dynamic connectivity (Dynamic network connectivity and saccadic behavior coupling). Static connectivity measures were used as predictors, and behavioral variability measures (CV-NormPV and CV-RT) as responses, with separate models estimated for HC and PD.

To examine the relative contributions of the Cognitive Saccade Network (CSN), Motor Saccade Network (MSN), and Cognitive-Motor Saccade Network (CMSN), PLS regression was refit separately within each sub-network and using only that specific network’s edges as predictors, as described for dynamic connectivity. The same within-group bootstrap and between-group permutation procedures described above were applied to each sub-network, with Holm correction applied across the three sub-networks, computed separately for each behavior and group.

## Supplementary results

### Saccadic task

All results on task performance and saccadic behavior have been presented in the main results section. Saccadic behavior changes with PD.

### Dynamic network connectivity and saccadic behavior coupling

Full statistics for the within-group and between-group whole-network analyses reported in the main text are provided in **Supplementary Table 2**. Within-group estimates derive from bootstrap resampling (5000 iterations); between-group estimates derive from subject-level group-label permutation testing (5000 iterations). Within-group and between-group p-values were each Holm-corrected within their own family of two behavioral comparisons (CV-NormPV, CV-RT), as described in Supplementary Methods.

**Supplementary Table 2.** Saccade network - behavior coupling, within-group and between-group statistics.

| Comparison | Group | Behavior | Mean estimate | 95% CI | p | p<sub>Holm</sub> |
| --- | --- | --- | --- | --- | --- | --- |
| Within-group (vs. zero) | HC | CV-NormPV | 0.00125 | [0.00068, 0.00181] | **0.0024** | **0.0048** |
| Within-group (vs. zero) | HC | CV-RT | 0.00010 | [-0.0006, 0.0008] | 0.7676 | 0.7676 |
| Within-group (vs. zero) | PD | CV-NormPV | - 0.00030 | [-0.0011, 0.0008] | 0.4904 | 0.5448 |
| Within-group (vs. zero) | PD | CV-RT | - 0.00049 | [-0.0012, 0.0006] | 0.2724 | 0.5448 |
| Between-group (HC vs. PD) | - | CV-NormPV | - | - | **0.0110** | **0.0220** |
| Between-group (HC vs. PD) | - | CV-RT | - | - | 0.1656 | 0.1656 |
| **Bold indicates comparisons that were significant ($p < 0.05$)** |  |  |  |  |  |  |

Full statistics for the within-group and between-group comparisons for the cognitive and motor network contributions to saccadic behavior that are reported in the main text are provided in **Supplementary Table 3**. Within-group estimates derive from bootstrap resampling (5,000 iterations), and between-group estimates derive from subject-level group-label permutation testing on models refit separately within each sub-network.

**Supplementary Table 3.** CSN, MSN and CMSN network behavior coupling, within-group and between-group statistics.

| Network | Comparison | Group | Behavior | Mean estimate | 95% CI | Raw p | Holm p |
| --- | --- | --- | --- | --- | --- | --- | --- |
| CSN | Within-group (vs. zero) | HC | CV-NormPV | 0.0058 | [0.0012, 0.0094] | **0.0200** | **0.0200** |
| CSN | Within-group (vs. zero) | PD | CV-NormPV | -0.0021 | [-0.0051, 0.0014] | 0.1952 | 0.5856 |
| CSN | Within-group (vs. zero) | HC | CV-RT | 0.0014 | [-0.0037, 0.0062] | 0.5960 | 1.0000 |
| CSN | Within-group (vs. zero) | PD | CV-RT | -0.0021 | [-0.0055, 0.0015] | 0.2864 | 0.4272 |
| CSN | Between-group (HC vs. PD) | - | CV-NormPV | - | - | **0.0056** | **0.0168** |
| CSN | Between-group (HC vs. PD) | - | CV-RT | - | - | 0.1334 | 0.4001 |
| MSN | Within-group (vs. zero) | HC | CV-NormPV | 0.0027 | [0.0009, 0.0043] | **0.0064** | **0.0168** |
| MSN | Within-group (vs. zero) | PD | CV-NormPV | -0.0003 | [-0.0021, 0.0016] | 0.8060 | 1.0000 |
| MSN | Within-group (vs. zero) | HC | CV-RT | -0.0010 | [-0.0031, 0.0010] | 0.3088 | 0.9264 |
| MSN | Within-group (vs. zero) | PD | CV-RT | -0.0011 | [-0.0026, 0.0007] | 0.2136 | 0.4272 |
| MSN | Between-group (HC vs. PD) | - | CV-NormPV | - | - | 0.0674 | 0.0674 |
| MSN | Between-group (HC vs. PD) | - | CV-RT | - | - | 0.5997 | 0.5997 |
| CMSN | Within-group (vs. zero) | HC | CV-NormPV | 0.0020 | [0.0006, 0.0030] | **0.0056** | **0.0168** |
| CMSN | Within-group (vs. zero) | PD | CV-NormPV | -0.0002 | [-0.0015, 0.0011] | 0.7580 | 1.0000 |
| CMSN | Within-group<br>(vs. zero) | HC | CV-RT | 0.0003 | [-0.0010,<br>0.0016] | 0.6176 | 1.0000 |
| CMSN | Within-group<br>(vs. zero) | PD | CV-RT | -0.0009 | [-0.0020,<br>0.0003] | 0.1228 | 0.3684 |
| CMSN | Between-group (HC<br>vs. PD) | - | CV-NormPV | - | - | **0.0288** | 0.0576 |
| CMSN | Between-group (HC<br>vs. PD) | - | CV-RT | - | - | 0.2054 | 0.4107 |
| ***Bold indicates comparisons that were statistically significant ($p < 0.05$)*** |  |  |  |  |  |  |  |

### Static connectivity

Group differences in static connectivity were largely consistent with those observed for dynamic connectivity in the main analysis (**Figure 2**). The group-averaged static connectivity across the whole saccade network was lower in PD than in HC. Using the same non-parametric label permutation approach (n = 10,000; threshold at the 95th percentile), group differences (HC minus PD) revealed a widespread reduction in connectivity across the saccade network (**Supplementary Figure 3**). Of the 741 edges, 515 edges (68.2%) showed significant differences between groups, with the vast majority (98.06%) exhibiting reduced connectivity in PD. As observed for dynamic connectivity, altered edges were distributed across all subnetworks, with the largest proportion involving the CSN (43.2%), followed by the CMSN (29.2%) and MSN (27.6%). Overall, these findings indicate that static connectivity reproduces the widespread network disintegration observed using dynamic connectivity measures.

**Supplementary Figure 3.**
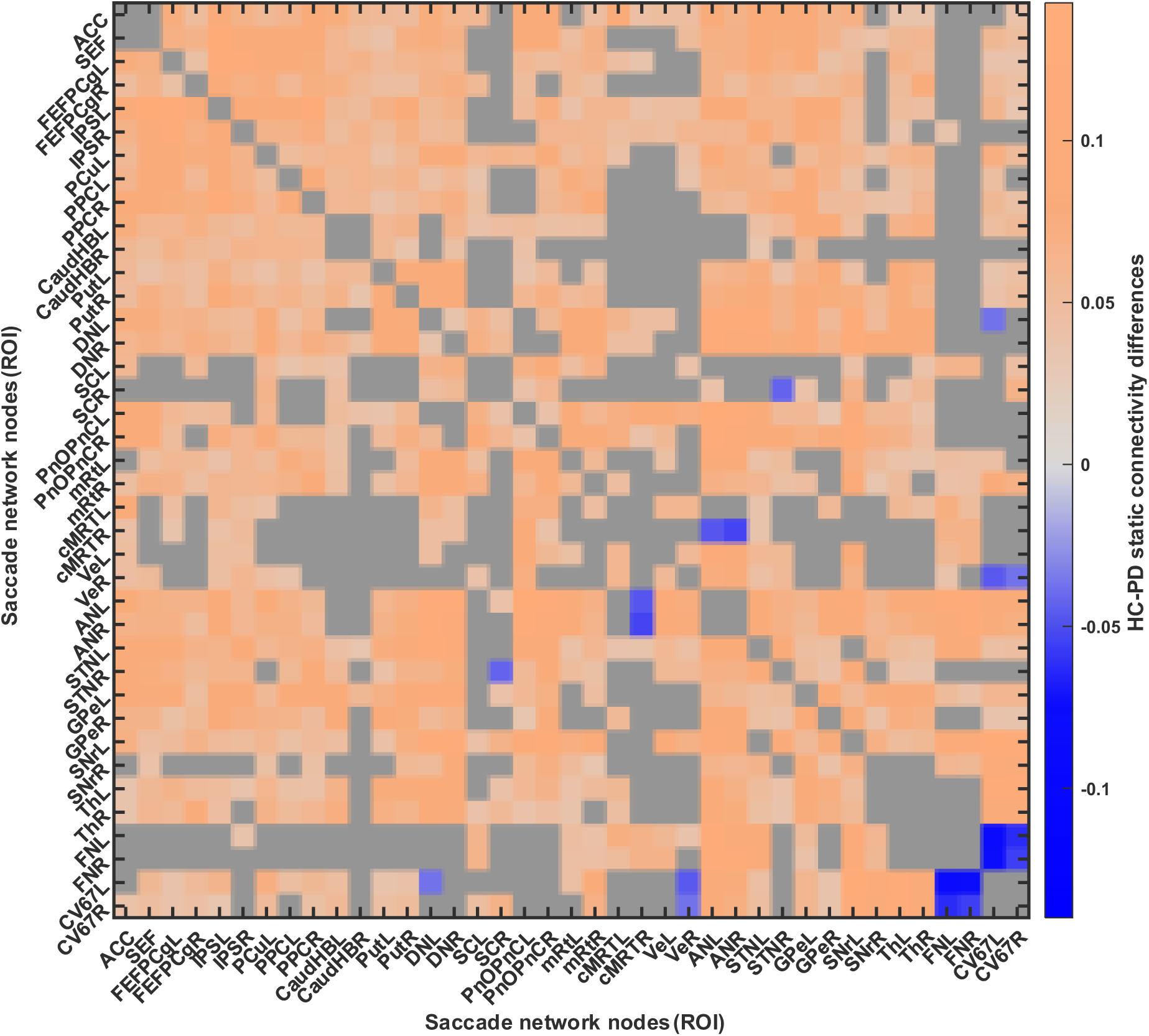
Static connectivity difference between HC and PD groups: Static connectivity matrix displaying the spread of group differences over the edges. The colorbar shows the actual differences in Pearson correlation coefficients, and the gray edges are the non-significant ones after permutation testing.

The partial least squares regression analysis used 741 connectivity values from the upper half of the static functional connectivity matrices to predict two behavioral measures (CV-NormPV and CV-RT) in HC and PD separately. Overall, results obtained with static connectivity (**Supplementary Figure 4**) were largely consistent with those from dynamic connectivity (**Figures 3 and 4**). Using static connectivity, whole saccade network coupling to CV-NormPV was reliably present in HC (p = 0.002, p_Holm_ = 0.005) but not in PD (p = 0.46, p_Holm_ = 0.60), with a significant between-group difference (p = 0.02, p_Holm_ = 0.04). CV-RT showed no reliable coupling in either group and no between-group difference (all p ≥ 0.22, p_Holm_ ≥ 0.22). In the cognitive and motor networks, HC showed reliable coupling to CV-NormPV in CSN (p = 0.01, pHolm = 0.01), MSN (p = 0.002, p_Holm_ = 0.005), and CMSN (p = 0.005, p_Holm_ = 0.01), which was absent in PD across all three (all p_Holm_ ≥ 0.34). The between-group difference in CV-NormPV coupling was significant only for CSN (p = 0.003, p_Holm_ = 0.01); MSN (p = 0.12, p_Holm_ = 0.12) and CMSN (p = 0.03, p_Holm_ = 0.05) showed non-significant trends. None of the CSN, MSN, or CMSN networks showed a significant relationship to CV-RT within either group or between groups (all p_Holm_ ≥ 0.68). Full statistics are provided in **Supplementary Table 4**.

**Supplementary Figure 4.**
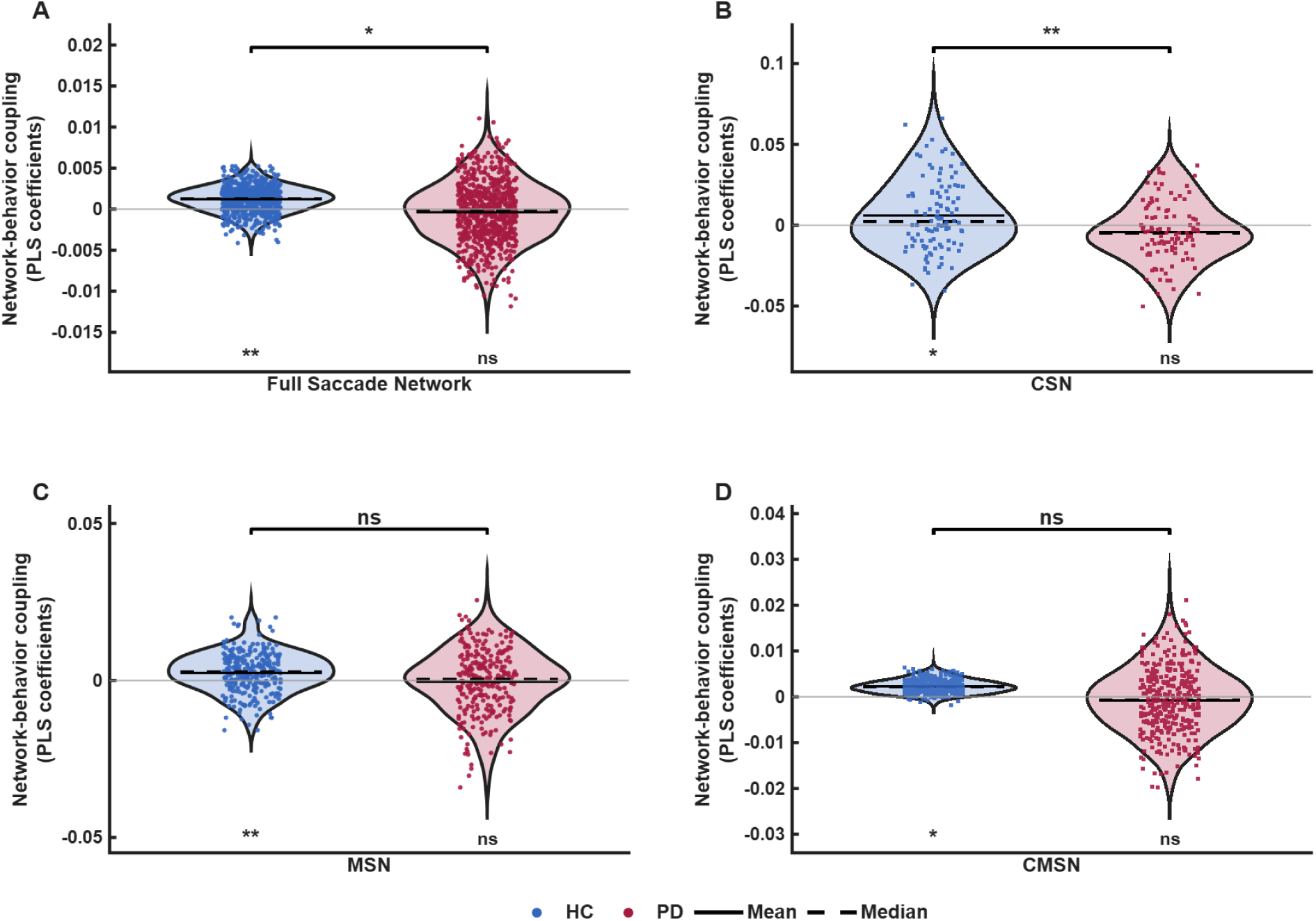
Static connectivity-based saccadic network-behavior coupling analysis: Distribution and comparison of PLS-derived network connectivity to saccadic variability coupling coefficients for healthy controls (HC, blue) and individuals with Parkinson’s disease (PD, red) obtained from static connectivity measures.Violin plots depict the distribution of edgewise coupling coefficients, with solid and dashed horizontal lines indicating the mean and median, respectively. Positive coefficients indicate that stronger connectivity is associated with greater behavioral variability, whereas negative coefficients indicate that stronger connectivity is associated with lower behavioral variability. Results are shown for the coefficient of variation of normalized peak velocity (CV-NormPV) for the full saccade network (A), CSN (B), MSN (C), and CMSN (D). Asterisks below each violin indicate whether the mean coupling coefficient differed significantly from zero within each group, while brackets denote between-group comparisons (*p < 0.05, **p < 0.01, ns-not significant).

**Supplemental Table 4.**
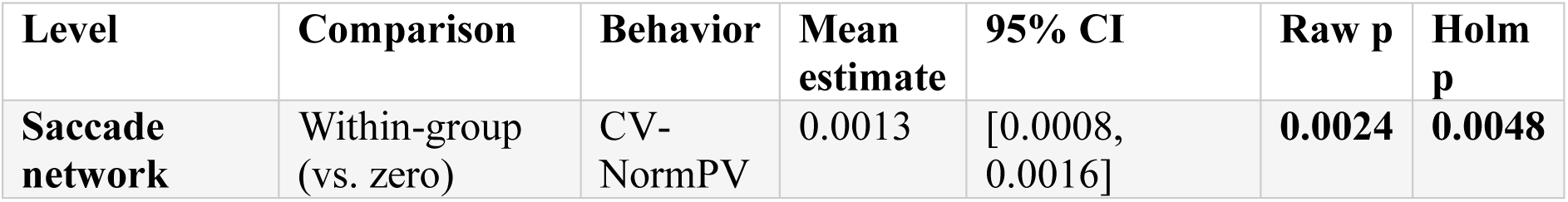
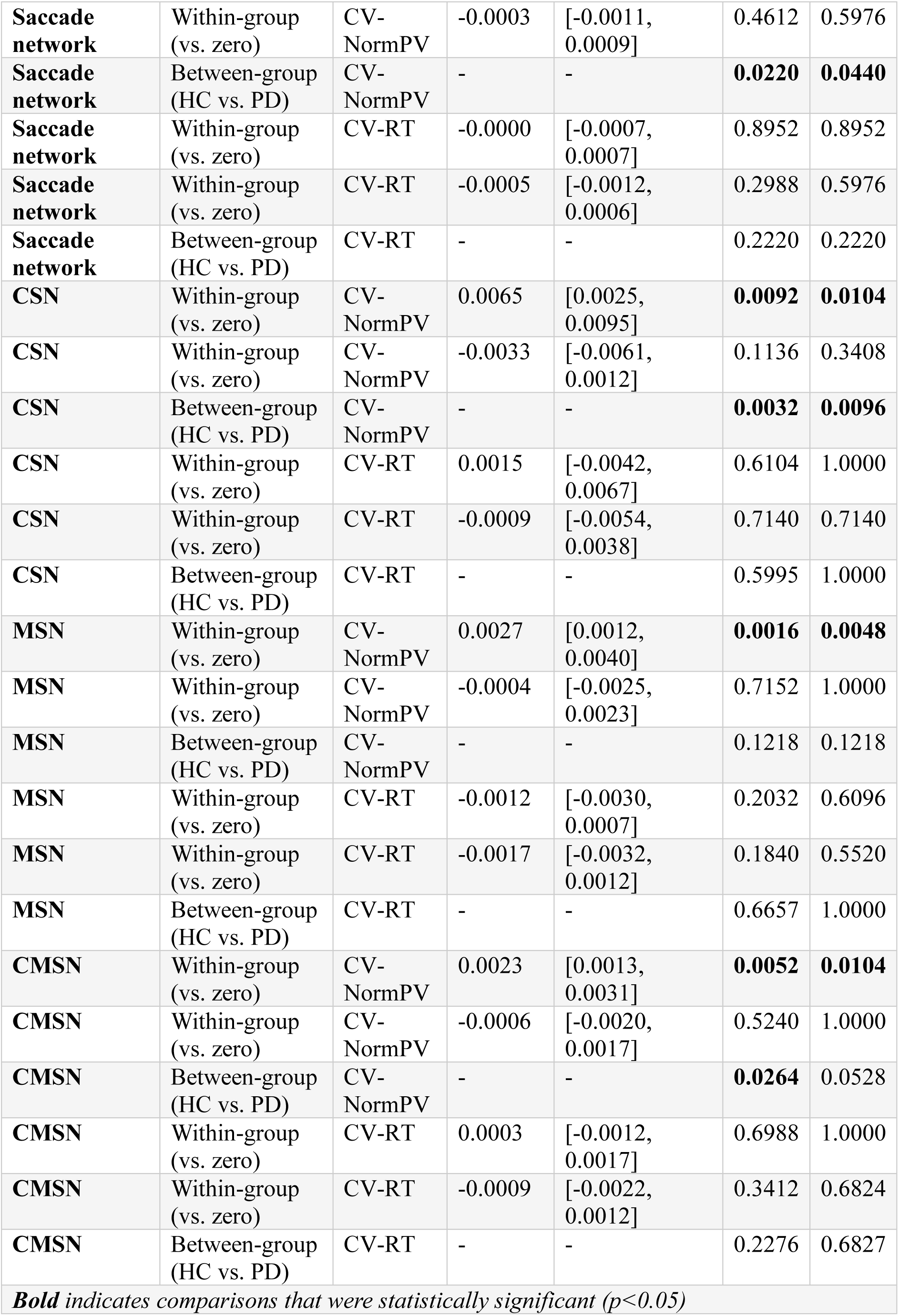
Static connectivity coupling: within-group and between-group statistics.

